# NickSeq for genome-wide strand-specific identification of DNA single-strand break sites with single nucleotide resolution

**DOI:** 10.1101/867937

**Authors:** Juniper J. Elacqua, Navpreet Ranu, Sarah E. Dilorio, Paul C. Blainey

**Affiliations:** MIT Department of Biological Engineering, Cambridge, MA, 02139, USA; Broad Institute of MIT and Harvard, Cambridge, MA, 02142, USA; Koch Institute for Integrative Cancer Research at MIT, Cambridge, MA, 02142, USA; 279 East Grand Avenue, South San Francisco, CA, 94080, USA; Department of Biomedical Engineering, Eindhoven University of Technology, 5600 MB Eindhoven, The Netherlands

## Abstract

DNA single-strand breaks (SSBs), or ‘nicks’, are the most common form of DNA damage. Nicks occur at rates of tens of thousands per cell per day, and result from many sources including oxidative stress and endogenous enzyme activities. Accumulation of nicks, due to high rates of occurrence or defects in repair enzymes, has been implicated in multiple diseases. However, improved methods for nick analysis are needed to learn how their locations and number affect cells, disease progression, and health outcomes. In addition to natural processes including DNA repair, leading genome-editing technologies rely on nuclease activity, including nick generation, at target sites. There is currently a pressing need for methods to study unintended nicking activity genome-wide to evaluate the impact of emerging genome editing tools on cells and organisms. Here we developed a new method, NickSeq, for efficient strand-specific profiling of nicks in complex DNA samples with single nucleotide resolution and low false-positive rates. NickSeq produces deep sequence datasets enriched for reads near nick sites and establishes a readily detectable mutational signal that allows for determination of the nick site and strand. In this work, we apply NickSeq to profile off-target activity of the Nb.BsmI nicking endonuclease and an engineered spCas9 nickase. NickSeq will be useful in exploring the relevance of spontaneously occurring or repair-induced DNA breaks in human disease, DNA breaks caused by DNA damaging agents including therapeutics, and the activity of engineered nucleases in genome editing and other biotechnological applications.

## INTRODUCTION

Cells continuously accrue DNA damage due to exposure to environmental stressors. The single-strand break (SSB), or ‘nick’, is estimated to occur at rates of tens of thousands per cell per day, making it the most frequent type of damage.^1^ Reactive oxygen species such as hydrogen peroxide-derived free radicals can nick DNA either directly by reacting with its sugar-phosphate backbone or indirectly by altering nucleobases which are then subject to repair pathways entailing nicked intermediates.^1,2^ Endogenous processes including DNA replication and topoisomerase activity also generate nicks.^1,3^

Accumulation of nicks, be it due to increased DNA damage, decreased DNA repair activity, or dysregulated endogenous processes, has many consequences. For example, DNA lesions, including nicks, can block RNA polymerase progression and result in incomplete transcription of genes that contain such lesions.^4^ Furthermore, nicked genomic DNA can lead to replication fork collapse during DNA synthesis, resulting in double-strand breaks (DSBs) and insertion/deletion (indel) mutations after error-prone DSB repair.^5^ Excessive damage can activate poly(ADP-ribose) polymerase (PARP), a protein capable of identifying nicks, signaling for their repair, and triggering cell death via necrosis or apoptosis.^6^ Nicks have also been implicated in heart failure in mouse models through increased expression of inflammatory cytokines^7^ and are thought to be a major factor affecting the rate of telomere shortening.^8,9^ Ataxia-oculomotor apraxia 1 (AOA1)^10^ and spinocerebellar ataxia with axonal neuropathy 1 (SCAN1)^11^ are neurodegenerative disorders associated with deficient repair of nicks resulting from abortive DNA ligase and topoisomerase 1 activity, respectively. Finally, deficient nick repair is observed in many tumors.^1^

In addition to the spontaneous accumulation of breaks, enzymes able to make engineered site-specific DNA breaks are fundamental to recombinant DNA technology and important genome editing technologies. RNA-guided nucleases, such as CRISPR associated protein 9 (Cas9) and its engineered nickase variants, represent simple and robust tools for targeted genome editing due to their ability to generate site-specific DNA DSBs and nicks in cells.^12–15^ Cas9 exhibits off-target activity, however, and the off-target DSBs the wild-type protein generates can lead to significant unintended mutagenesis, toxicity, and cell death.^16,17^ In efforts to make precise edits and minimize off-target DSB toxicity, Cas9’s engineered nickase variants are an increasing focus in genome editing applications.^18–25^

In practice, two guide RNAs can be designed to flank a target site and generate nicks on opposite DNA strands that together compose an on-target DSB with reduced off-target DSBs.^19^ CRISPR base editors are composed of Cas9 nickase enzymes fused to either a cytidine deaminase or an adenosine deaminase enzyme to generate targeted C→T or A→G mutations, respectively.^20,21^ The nick is incorporated on the opposite DNA strand of the desired mutation to increase editing efficiency by stimulating mismatch repair at the site before base excision repair can replace the deaminated nucleotide.^20,21^ Additionally, it has been shown that insertions can be made at nicks via homology directed repair with a designed donor DNA without non-homologous end joining (NHEJ) and the potential for undesired indel mutations.^22^ Creating these nicks with Cas9 nickase can also expand the targeting scope for editing, as desired edits can reside outside of the guide RNA target sequence and PAM.^23,24^ Furthermore, repair at nicks shows significantly higher efficacy than repair at DSBs, measured as ratio of desired editing to unintended mutagenesis such as indels formed from NHEJ.^22^ Finally, the recently reported prime editing strategy can generate specific base substitutions, insertions, and deletions by initiating DNA synthesis at a precisely located nick using an extended guide RNA as the template for reverse transcription.^25^

While nicks are less toxic and pose less potential for unwanted indel generation than DSBs, the off-target activity of Cas9 nickase remains a concern. Large numbers of nicks can still interrupt cellular functions and lead to cell death^4–6^ and it has been shown that nicking base editors can cause off-target modifications.^26^ Furthermore, homology directed repair at off-target nicks can lead to loss of heterozygosity, a form of genetic instability commonly observed in tumor cells.^24^ The potential for unintended, potentially oncogenic modifications in even a rare subset of cells is a significant concern for clinical applications of genome editing. Genome editing technologies that rely on Cas9 nickases are not commonly tested for off-target nicking activity directly, as high-resolution nick detection technologies have not been available. Rather, off-target activity is predicted by proxy by analysis of off-target DSB sites of the nickases’ wild-type Cas9 counterparts.^20,21,25^

Available methods for direct, highly resolved assessment of nicks are lacking, which has limited the analysis of off-target nicks in genome editing. The alkaline comet assay reveals the overall extent of nicks and other alkali-labile sites (such as abasic sites) when compared with the results of a neutral comet assay, but does not provide information about the locations of breaks.^27^ The SSB-seq method was developed to map nicks to the genome through nick translation with digoxigenin-modified nucleotides followed by anti-digoxigenin immunoprecipitation, although it does not achieve single nucleotide resolution.^28^ While preparing this manuscript, we became aware of a new high-performance nick detection method termed Nick-seq that works using a different procedure and was applied for the detection of processed DNA lesions in bacterial samples.^29^

Here, we built upon SSB-seq^28^ to create a method capable of identifying DNA nicks with single nucleotide resolution in human genomic DNA. This method, which we call NickSeq, relies on an engineered mutational signal arising from nick translation with degenerate nucleotides.^30^ Utilizing this signal in conjunction with the enrichment of reads near nicks allows us to achieve single nucleotide resolution and strand-specific detection of nicks with low false-positive rates. We used NickSeq to assess the off-target activities of the nicking endonuclease Nb.BsmI and the D10A nickase variant of spCas9. NickSeq has many potential applications that have been underserved by previous methods for nick detection. Like Nick-seq, NickSeq could also be used to study the activity of DNA damaging agents such as reactive oxygen species and to explore in quantitative detail the etiology of AOA1^10^ and SCAN1,^11^ two neurodegenerative disorders characterized by deficient nick repair.

## RESULTS

### Degenerate nucleotides provide a mutational signal demarking the location of nicks

Our approach takes advantage of synthetic non-canonical deoxynucleotides capable of base pairing with more than one of the four naturally occurring deoxynucleotides. The bases 6*H*,8*H*-3,4-dihydropyrimido[4,5-*c*][1,2]oxazin-7-one (P) and *N*^6^-methoxy-2,6-diaminopurine (K) act as a universal pyrimidine and purine, respectively (Fig. 1a).^30^ dPTP and dKTP can be incorporated into a DNA molecule at the site of a nick via nick translation (Fig. 1b) and generate readily detectable sequence variants when amplified by PCR (Fig. 1c). To characterize the mutational signatures that arise from P and K, we annealed custom oligonucleotides to obtain four DNA molecules, each with a nick present just 5’ of each of the four canonical deoxynucleotides (Fig. 2a). After PCR with *Taq* DNA polymerase and sequencing, we found evidence of P and K incorporation extending 4-8 bases downstream of the original nick site, with the ratio of C:T at sites where P was incorporated being ∼40:60 and the ratio of G:A at sites where K was incorporated being ∼15:85 in agreement with a previous study (Fig. 2b).^30^ PCR with KAPA HiFi DNA polymerase also results in 4-8 downstream bases displaying evidence of dPTP and dKTP incorporation, though with different distributions of C:T and G:A at such sites (Fig. 2c). We considered other degenerate nucleotides for use in NickSeq, such as inosine (I) and ribavirin (R). I is recognized as G by DNA polymerases during PCR,^31^ however, and as such would not allow for true single nucleotide resolution of nicks flanked by G residues. R was considered in place of K as a universal purine^32^ but produced less reliable results (Fig. S1).

**Figure 1:**
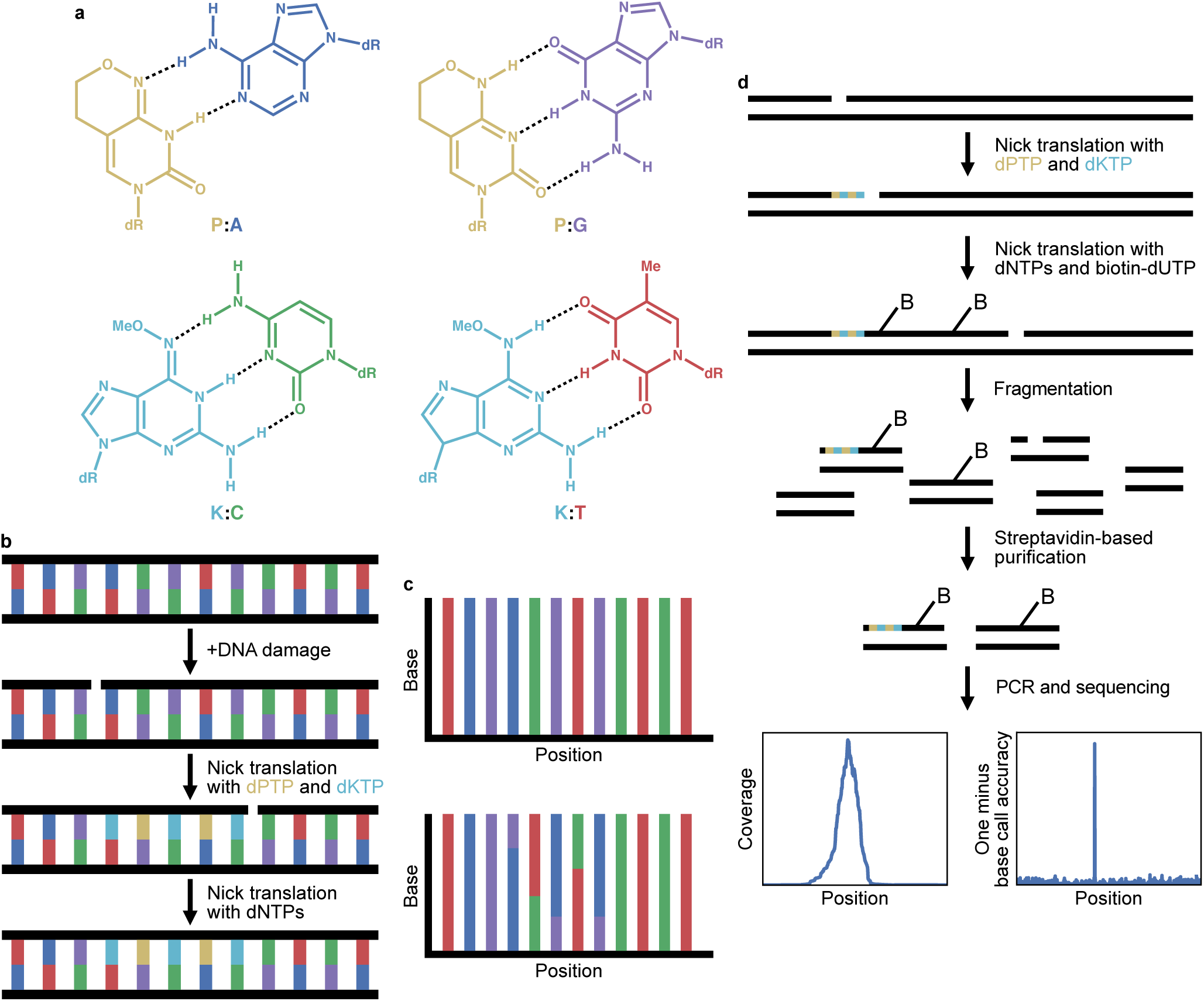
NickSeq overview and workflow. **a)** Chemical structures of the two degenerate nucleotides dPTP and dKTP in both of their tautomeric forms enabling their respective activities as universal pyrimidine and purine. Black dotted lines represent hydrogen bonds between nucleotides. **b)** During nick translation with only the two degenerate nucleotides, P residues (gold) are inserted across from A (blue) and G (purple) residues while K residues (cyan) are inserted across from C (green) and T (red) residues. **c)** Sequencing DNA fragments without nicks or that had nick translation occur with only regular dNTPs will not show a mutational signal (top). Sequencing DNA fragments that underwent nick translation with dPTP and dKTP will show a mutational signal that extends a few bases 3’ of the nick’s original location (bottom). **d)** Workflow used to perform NickSeq. Nick translations are performed consecutively with dPTP plus dKTP and then with regular dNTPs plus biotinylated dUTP. Of note, nick translation will move a nick from its original location but will not actually repair the nick. DNA is fragmented and streptavidin-based purification enriches for fragments around the original (pre-nick translation) sites of nicks. After PCR and sequencing, sequence coverage and mutational information allow for sensitive single nucleotide-resolved and strand-specific identification of nick sites.

**Figure 2:**
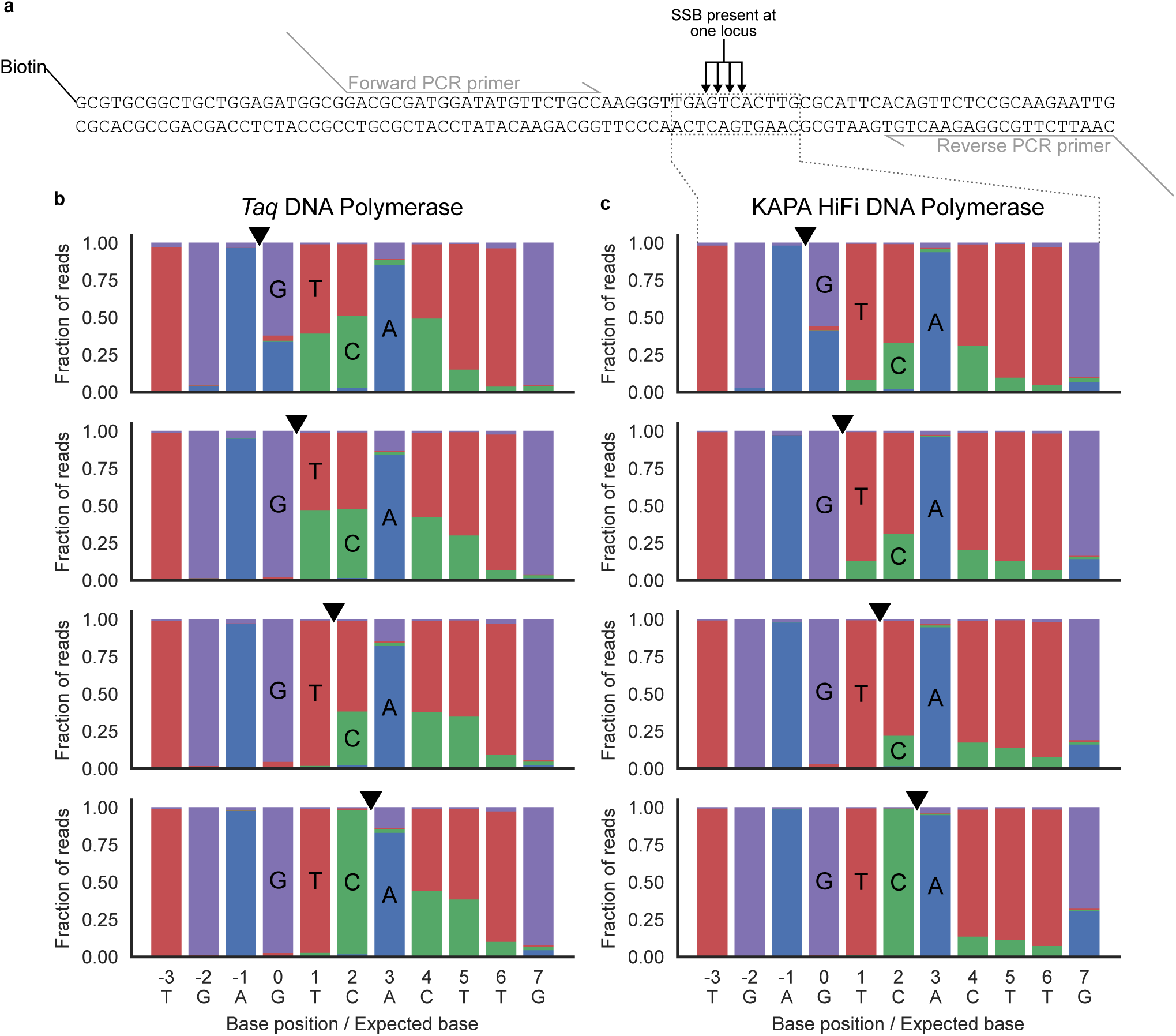
Effect of degenerate nucleotides during PCR. **a)** Schematic of the oligonucleotides used to test the mutational outcome when dPTP and dKTP are incorporated into DNA and amplified by PCR. Four different oligonucleotides were generated, each with a nick directly 5’ of a different one of the four native dNTPs. The strand with the nick also contained a 5’ biotin-TEG modification to allow for purification of just that strand prior to PCR. PCR primers were designed to specifically amplify the region of the oligonucleotides containing the nicks and were tailed with 5’ sequences compatible with secondary PCR using barcoded P5 and P7 sequencing primers. **b and c)** Sequencing result of the four oligonucleotides following consecutive nick translations with dPTP plus dKTP and then with standard dNTPs, purification, and PCR with *Taq* DNA polymerase **(b)** or KAPA HiFi DNA polymerase **(c)**. The black triangles represent the locations of the nicks for each sample.

### Biotinylated nick translation products can be enriched to focus sequencing effort at nick sites and enable confident strand determination

Nick translation of biotin tagged nucleotides after incorporation of the degenerate nucleotides allows for selective pulldown of DNA molecules containing P and K residues via a streptavidin-based purification. Increased sequencing coverage around nick sites provides independent evidence of nicks that can be analyzed in conjunction with the mutational signal (Fig. 1d). To test our library enrichment, we digested a plasmid containing two recognition sites with the nicking endonuclease Nb.BsmI (Fig. 3a). After performing NickSeq on this sample, we observed up to 1,000 fold higher coverage of sequence reads at sites around the nicks (Fig. 3b). These coverage peaks can be identified by MACS2, a peak caller originally designed for use with ChIP-seq data,^33^ while the mutational signal from P and K incorporation reveal the exact location of nicks within the peaks (Fig. 3c,d). These signals are robust to the enzyme used to incorporate the biotinylated nucleotides (Fig. S2) and the enzyme used to conduct PCR (Fig. S3). Furthermore, similar levels of enrichment are achieved when using nucleotides that are modified with desthiobiotin (Fig. S4).^34^ Peak width and the magnitude of enrichment are highly dependent on the concentration of nucleotides during the nick translation reaction (Fig. S5).

**Figure 3:**
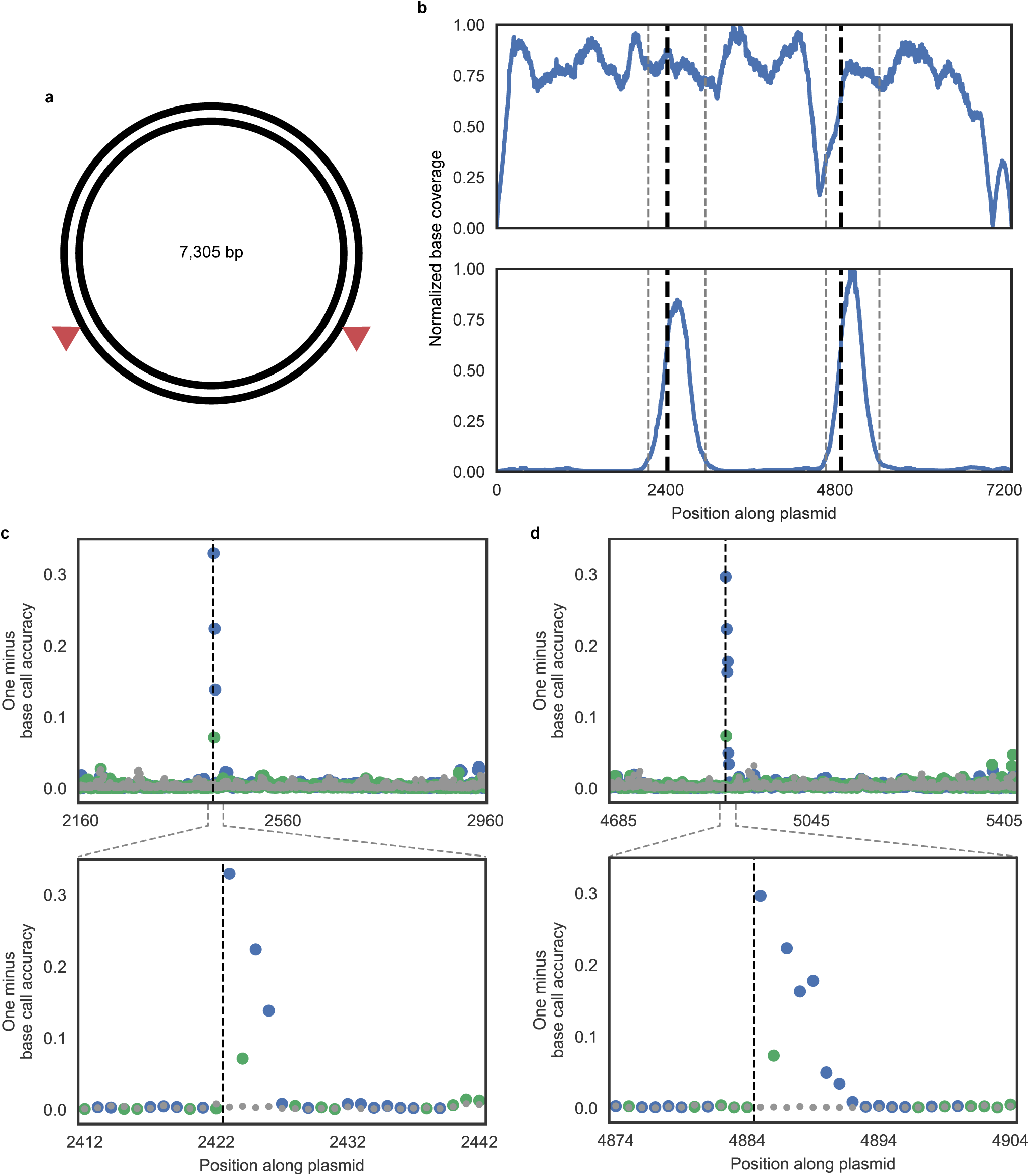
Single base pair resolution of nicks in plasmid DNA. **a)** Schematic of the plasmid used to test NickSeq where red arrows represent the expected locations of nicks after treatment with the nicking endonuclease Nb.BsmI (both are expected on the reference strand). **b)** Sequencing coverage normalized to the location with maximum read depth for untreated plasmids (top) and with two nicks introduced by Nb.BsmI treatment (bottom) after performing NickSeq. Black dashed lines represent the expected locations of nicks after Nb.BsmI treatment and gray dashed lines represent the bounds of MACS2 peaks called on data from the treated sample. **c and d)** Mutational signal for all locations within the called MACS2 peaks (top) and close-up views where there is high mutational signal (bottom). Black dashed lines represent the expected locations of nicks after Nb.BsmI treatment. Data in gray represent the untreated plasmid while data in blue and green represent the Nb.BsmI treated sample where a P or K residue would be inserted after a reference-strand nick, respectively.

Library enrichment also plays a crucial role in determining the strandedness of a nick. As DNA polymerases extend DNA in the 5’→3’ direction and the nick translation steps are carried out consecutively, the biotinylated nucleotides will always be 3’ of the degenerate nucleotides and the position with maximum sequencing coverage occurs 3’ of the mutational signal. Sequencing data is typically viewed with respect to the reference strand of double stranded DNA, so a nick that occurred on the reference strand will result in the coverage peak appearing 3’ of the mutational signal. A nick that occurred on the non-reference strand, however, will result in the coverage peak appearing 5’ of the mutational signal since it is on the 3’ side with respect to the non-reference strand (Fig. S6). Determining the strandedness of a nick is, in turn, important for single nucleotide resolution since multiple P and K residues can be incorporated at a nick resulting in a few consecutive bases with mutational signal. The nick is always just 5’ of the terminal position with mutational signal.

### Nicking endonuclease Nb.BsmI has detectable off-target activity

Nicking reactions set up with Nb.BsmI that contain too high an enzyme concentration, that proceed for too long, or that have slightly altered buffer environments result in off-target activity that is detectable by NickSeq (Fig. 4a). Enrichment of the two on-target sites still occurs, but to a lesser extent than when off-target nicks are absent. Since read coverage is relatively high throughout the plasmid, we did not restrict our nick calls to MACS2 peaks in this instance. In addition to the on-target sites, there are twelve sites with mutational signal above background level (Fig. 4b, Table S1). Each of these sites has a single nucleotide mismatch from the six nucleotide consensus Nb.BsmI recognition sequence, suggesting that they represent off-target activity of the enzyme. There are a total of 57 such sites in the plasmid, so twelve sites with high mutational signal occurring at such near-cognate Nb.BsmI sites by chance is highly unlikely (*p* < 10^−25^, hypergeometric test), suggesting these mutational signals indeed reflect the presence of nicks resulting from off-target Nb.BsmI activity.

**Figure 4:**
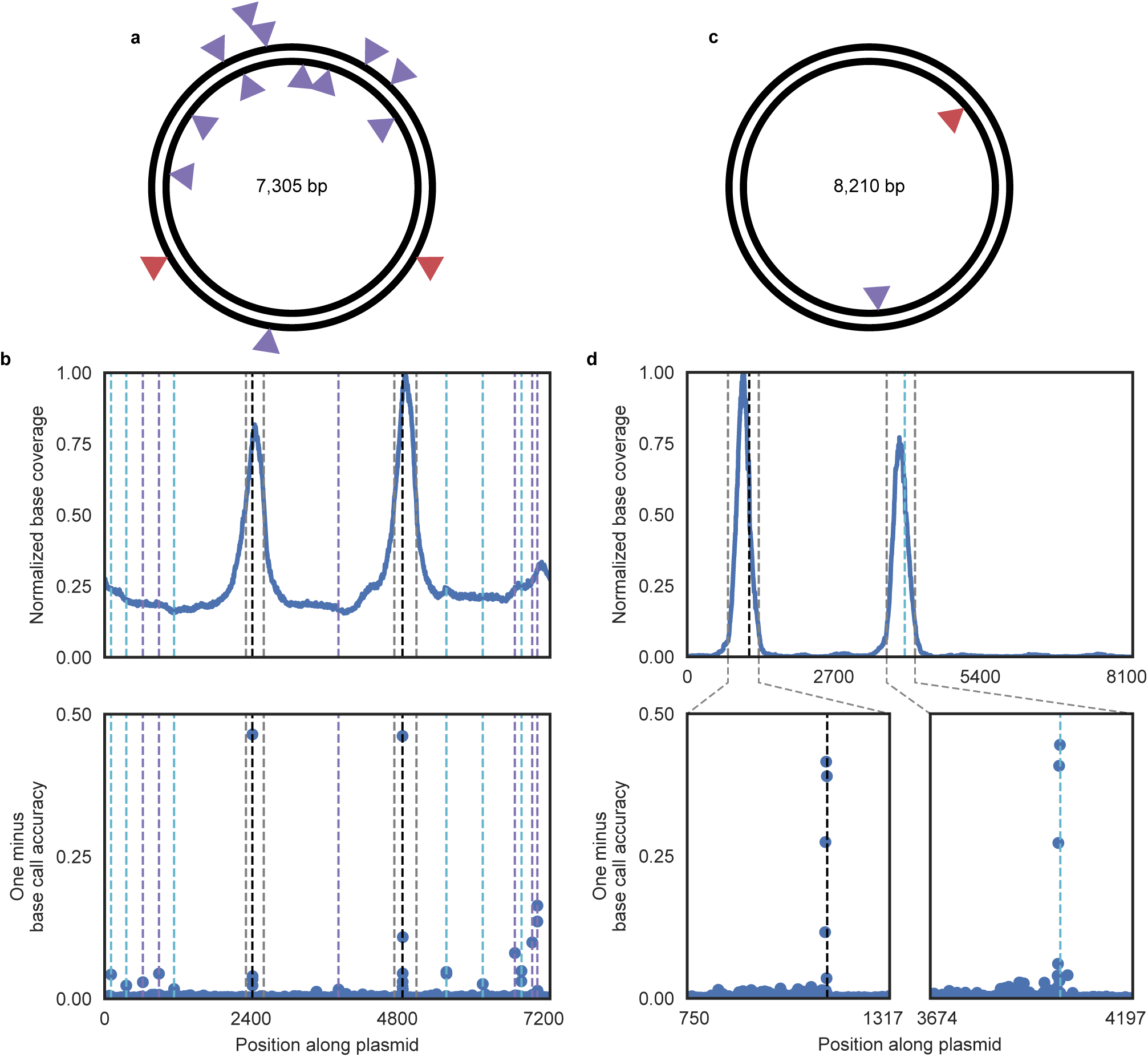
Detection of off-target nicks in plasmid DNA. **a)** Schematic of a plasmid used to test NickSeq where red arrows represent on-target activity of Nb.BsmI and purple arrows represent detected off-target activity of Nb.BsmI. **b)** Normalized sequencing coverage (top) and base call error rate (bottom) for the plasmid in **a**. Black dashed lines represent on-target locations of Nb.BsmI activity and gray dashed lines represent the bounds of called MACS2 peaks. Purple and cyan dashed lines represent detected off-target locations of Nb.BsmI activity on the reference and non-reference strands, respectively. **c)** Schematic of a plasmid used to test NickSeq where the red arrow represents on-target activity of D10A spCas9 nickase and the purple arrow represents detected off-target activity (both are on the non-reference strand). **d)** Normalized sequencing coverage (top) and base call error rate (bottom) for the plasmid in **c**. The black dashed line represents on-target activity of D10A spCas9 nickase while the cyan dashed line represents off-target activity. The gray dashed lines represent the bounds of called MACS2 peaks.

### Cas9 nickase has detectable off-target activity in plasmid DNA

Cas9 off-target activity has been observed at locations with significant, but not full, complementarity to the guide RNA.^35–39^ We were able to observe this phenomenon directly in D10A spCas9 nickase activity via NickSeq in plasmid DNA. We used a guide RNA with one fully complementary on-target site and one predicted off-target site where proper base pairing would occur for 15 of the 20 nucleotides of the guide RNA (Fig. 4c). Significant enrichment and a strong mutational signal occurred at both sites, with slightly lower enrichment observed at the off-target site (Fig. 4d, Table S2).

### Nick detection is possible at genome scale

After demonstrating the capabilities of NickSeq on plasmid DNA, our next goal was to detect nicks in a genome. We used D10A spCas9 nickase to incorporate nicks in an *E. coli* genome of 4,622,358 bp by carrying out a reaction utilizing two guide RNAs wherein we predicted nine nicks would occur at fully cognate target sites (Fig. 5a). After performing NickSeq, all nine nicks were identified with no false positive identifications. Eight nicks, all originating from the same guide RNA, resulted in highly significant sequencing coverage peaks (Fig. 5b) and single nucleotide resolution achieved using the mutational signal (Fig. 5c, S7a). The remaining nick, originating from the other guide RNA, was located within a MACS2 called peak but with lower enrichment. In fact, this peak could not be distinguished from background reads based on sequence coverage or MACS2 statistics alone (Fig. 5b). The mutational signal in this peak allowed the nick to be detected (Fig. 5d). We hypothesize that the difference in sequencing coverage between the peaks containing nicks is due to a difference in the levels of activity between the two guide RNAs causing differential penetrance of the resultant nicks. Other sequence coverage peaks either lacked mutational signal altogether, contained singleton mutational signal (not at multiple consecutive or near-consecutive nucleotides), or contained mutation types other than the transition mutations that dPTP and dKTP are known to cause (Fig. S7b). These signals likely arose from sources other than P and K incorporation, such as misincorporation of canonical dNTPs during PCR or sequencing errors, and as such were straightforwardly filtered out. We did not expect to observe off-target activity from either guide RNA in this experiment. One guide exhibited low on-target activity and there were no sites in the genome with significant homology to the other, high activity guide that lacked mismatches at the 3’ end of the target sequence and were also adjacent to the Cas9 PAM.

**Figure 5:**
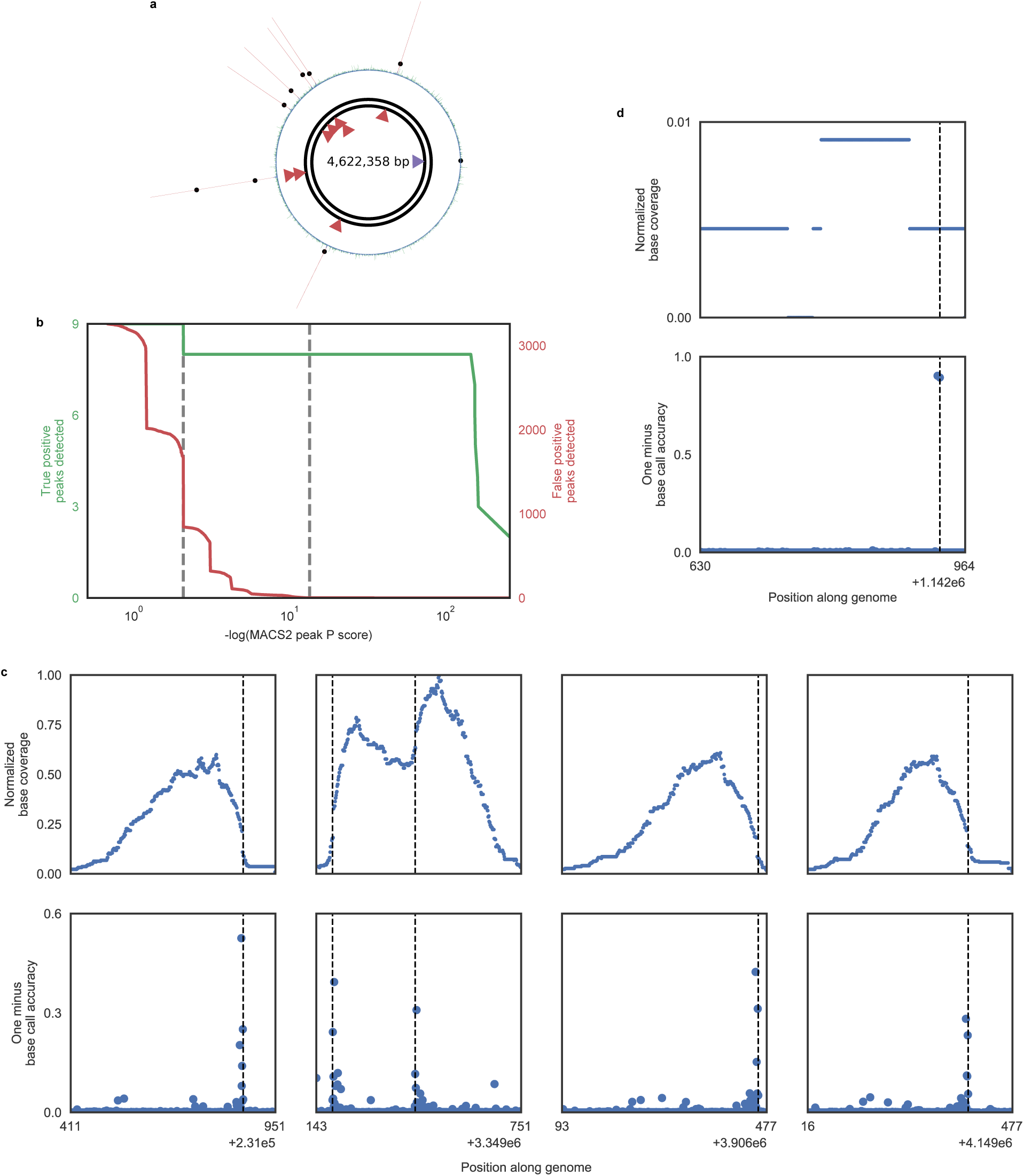
Single base pair resolution detection of nicks in genomic DNA. **a)** Schematic of an *E. coli* genome used to test NickSeq where red arrows represent the activity of Cas9 nickase paired with one guide RNA and the purple arrow represents the activity of Cas9 nickase paired with a different guide RNA. Circularized plot data show normalized sequencing coverage with regions in green representing the locations of called MACS2 peaks with quality score greater than 0.4. Lines in red represent locations of such peaks wherein a nick is identified with the exact location of that nick represented by a black dot. **b)** Cumulative number of nicks contained within MACS2 peaks (green) and cumulative number of MACS2 peaks containing no nick (red) as a function of p-value threshold. The left gray dashed line represents the p-value where the final nick-containing peak is identified and the right gray dashed line represents the p-value where the first peak not containing a nick is identified. This panel uses only the coverage signal and does not incorporate mutational data. **c)** Normalized sequencing coverage (top) and base call error rate (bottom) for four MACS2 peaks which contain nicks resulting from Cas9 nickase in conjunction with one of the guide RNAs used (corresponding to the red arrows in **a**). The second set of plots from the left represent a single MACS2 peak which encompassed two nicks due to their close proximity. **d)** Normalized sequencing coverage (top) and base call error rate (bottom) for the MACS2 peak that contains the nick resulting from Cas9 nickase in conjunction with the other guide RNA used (corresponding to the purple arrow in **a**). Coverage is lower for this guide, but there are still nucleotide positions proximal to the Cas9 nickase target site with high mutational signal that enable nick identification.

### Off-target Cas9 nickase activity can be detected across the human genome

We performed NickSeq on human genomic DNA treated with D10A spCas9 nickase and a guide RNA targeting the AAVS1 locus on chromosome 19 (which we also used in the plasmid experiment described above). This allowed for the identification of six nicks without using *a priori* knowledge of expected Cas9 on- or off-target activity to inform nick calling. Of these six nicks, the one with the highest enrichment is located at the on-target site (Fig. 6a) and the other five are located at off-target sites each with three or four mismatches to the guide RNA (Fig. 6b).

**Figure 6:**
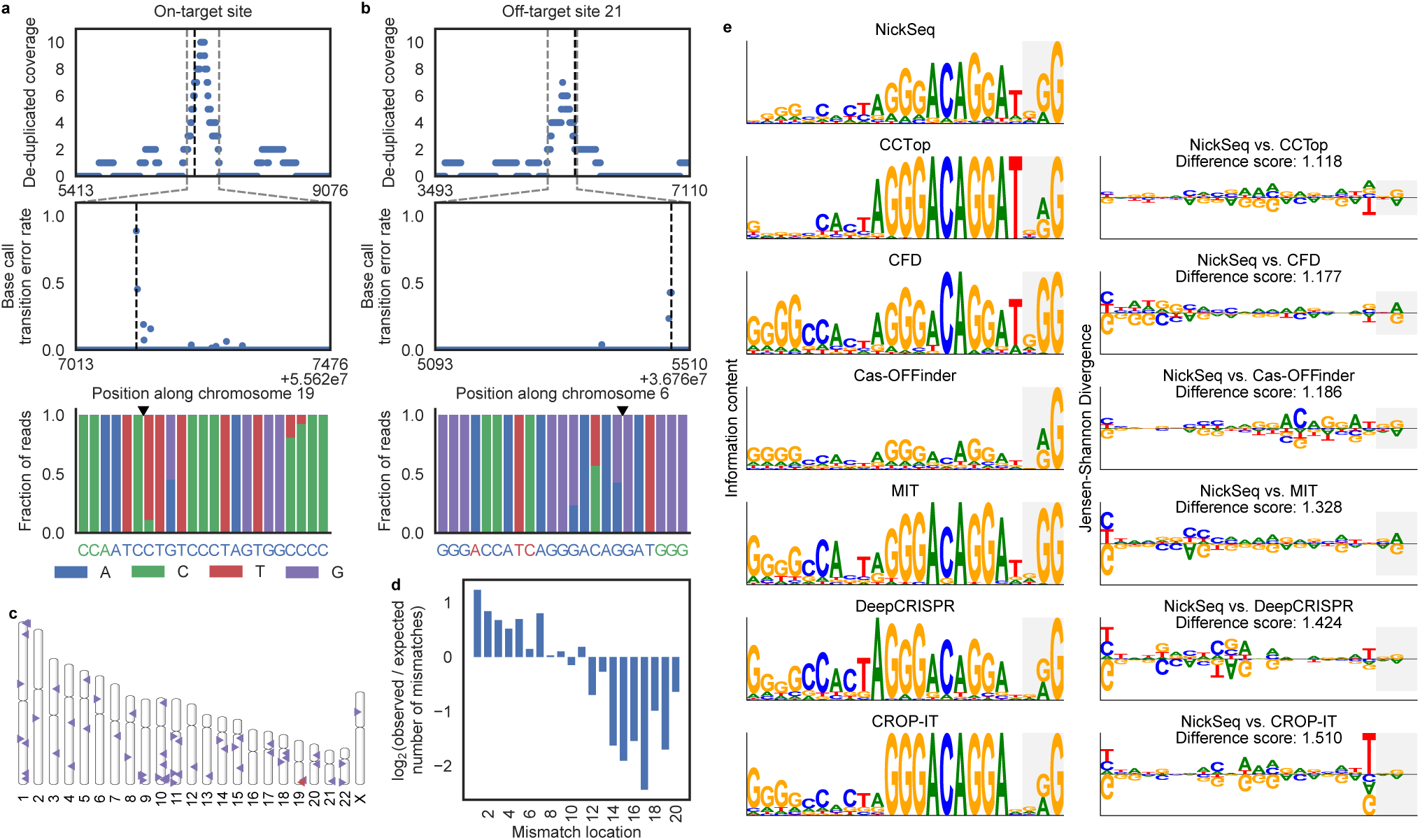
Detection and analysis of D10A spCas9 nickase on- and off-target activity in human genomic DNA. **a)** D10A spCas9 nickase single-site on-target activity is represented by the most prominent MACS2 peak, denoted by the gray dashed lines, which also contains strong mutation signal. The reverse complement of the guide sequence is shown in blue with the PAM in green at bottom. **b)** One of the five off-target nicks that were identified without utilizing *a priori* knowledge of expected off-target sites. Gray lines mark the MACS2 peak bounds. At bottom, the genomic sequence where the guide RNA binds is shown in blue with mismatches in red. **c)** Ideogram showing the site of on-target activity in red and candidate off-target sites with NickSeq mutational signal in purple. **d)** Comparison of the frequency of mismatch locations in the guide RNA at the sites of detected off-target activity to all potential sites proposed by Cas-OFFinder. **e)** Sequence motifs created from the sites with evidence of D10A spCas9 nickase off-target activity and the top hits predicted by multiple Cas9 off-target predictors (left). The motif for the sites identified by NickSeq is compared to the motifs for each of the software predictions using DiffLogo (right). The bases composing the PAM sequence are shown with a light gray background.

Nick calling can be enhanced further by considering only prospective off-target locations. To this end, we utilized Cas-OFFinder^40^ to identify all PAM-adjacent sites in the genome with up to eight mismatches to the guide RNA or with up to two mismatches and a two nucleotide DNA or RNA bulge. Analyzing these locations for sequence reads and mutational signal enabled the identification of 53 sites with evidence for off-target activity (Fig. 6c, Table S3). These off-target sites generally followed rules established to describe the off-target DSB activity of wild-type Cas9.^36^ Mismatches closer to the 5’ end of the guide RNA, distal from the PAM, are better tolerated than mismatches at the 3’ end, proximal to the PAM (Fig. 6d). Further, the PAM sequence 5’-NGG is the most prominent but there is observable off-target nicking activity at the alternative PAM 5’-NAG.

The guide RNA used here was previously tested for off-target activity in the human genome with wild-type spCas9.^41,42^ The methods previously used for DSB analysis with this particular guide were neither unbiased nor genome-wide, but rather relied on PCR and deep sequencing of select locations in the genome predicted as likely off-target sites, which hinders direct comparison to our findings. However, NickSeq provides evidence of D10A spCas9 nickase activity at five off-target sites where the previous studies described no wild-type spCas9 DSB off-target activity using the same guide RNA. Three of these five sites were detectable as nicks without the use of *a priori* knowledge of expected off-target sites and showed highly significant peak enrichment as measured by MACS2 (Table S3). Furthermore, the previous studies identified eight off-target sites with wild-type spCas9 DSB activity where NickSeq did not show D10A spCas9 nickase activity, suggesting that wild-type Cas9 and the D10A nickase mutant could have different off-target spectra.

We generated a sequence motif from the 52 sites of identified D10A spCas9 nickase activity where base-pairing between the guide RNA and target DNA did not result in a DNA or RNA bulge. In order to compare the D10A spCas9 nickase off-targets detected by NickSeq with predicted wild-type spCas9 off-targets we utilized several DSB off-target prediction tools. CCTop,^43^ CRISPOR,^44^ DeepCRISPR,^45^ and CROP-IT^46^ ranked all genome-wide potential off-targets and we generated sequence motifs for the top 52 sites proposed by each algorithm (for direct comparison with the 52 NickSeq-identified sites). Of note, two motifs were created based on CRISPOR’s output as it reports both MIT off-target scores (score calculation is based on data from Hsu, et al. (2013)^47^) and CFD off-target scores^48^ (Fig. 6e, left column). To compare the motifs created from our results and from each of the prediction tools, we utilized the DiffLogo strategy.^49^ Each position in the DiffLogo motif summarizes the base magnitude and polarity of difference between the two motifs being compared and enables calculation of a scalar total difference score between the motifs. We created DiffLogo motifs to perform pairwise comparisons between the NickSeq-identified off-targets and the proposed off-targets from each prediction tool, including the background distribution of all possible off-targets identified by Cas-OFFinder (Fig. 6e, right column). From the DiffLogo scores, the motif generated by the CCTop predictions most closely resembles the motif generated by the NickSeq results. The biggest discrepancy between these two motifs is at the base just next to the PAM region where the T is not as frequent at off-target sites as CCTop predicts. This is in contrast to CROP-IT and, to a lesser extent, DeepCRISPR where too little emphasis is placed on this position. For the CFD and MIT scores, it appears as though the bases at the 5’ end of the target sequence are being weighed too heavily and the 5’-NAG alternative PAM is nearly ignored.

All of the prediction tools utilized here were created from data collected with wild-type spCas9 and are meant to predict wild-type spCas9 DSB off-target activity. We expect there to be some differences in the off-targets created by wild-type spCas9 and its D10A nickase variant, so our analysis here is not a test of how well these tools perform at the task they were designed for. Rather, it reveals which currently available method for a related task performs best at predicting D10A spCas9 nickase off-target activity. As additional D10A spCas9 nickase off-targets are characterized, optimal parameters for each algorithm can be identified for predicting D10A spCas9 nickase activity and the prediction tools could have the option to search for expected wild-type/DSB or D10A nickase off-target activity. This could also be done with other spCas9 variants, such as the H840A nickase mutant, which may exhibit distinct off-target characteristics.

## DISCUSSION

Our results demonstrate a method for identifying nicks in human genomic DNA with single nucleotide resolution and strand specificity using a sequence variation signal. This technique relies on incorporating degenerate nucleotides at nicks through nick translation and reading out a mutational signal through sequencing. As this results in multiple consecutive transition mutations being identified, true signal can be readily distinguished from PCR misincorporations, sequencing errors, and other sources of noise which occur randomly and are unlikely to be present at consecutive nucleotides. In addition, library enrichment around nick sites reduces the sequencing requirement for nick detection in complex samples like large genomes. Further, the nicked strand can be identified by comparing the exact locations of the mutational signal and the peak in enriched sequence coverage.

We demonstrate the ability of NickSeq to identify nicks that should be rare within a sample of DNA molecules by detecting off-target activity of the nicking endonuclease Nb.BsmI and the D10A nickase variant of spCas9. Off-targets of these and related enzymes are important to quantify as they critically limit some applications including therapeutic genome editing. CRISPR/Cas9 nickase is used as an essential component of several advanced genome editing strategies.^19–25^ However, no method has been available to directly test the off-target activity of Cas9 nickase since nicks could not previously be detected with high resolution. Our results suggest that prior work to understand the off-target characteristics of wild-type Cas9 may not fully translate to nickases, underscoring the need for express analysis of off-target nicking to model off-target activity and support the development of next-generation nickase enzymes.

In addition to their significance in genome engineering, nicks occur in normal cellular functions and are relevant to human health and disease.^7–11^ Accumulation of nicks can lead to both apoptotic and necrotic cell death^6^ as well as dysregulation of cellular processes including transcription and DNA replication.^1^ Despite their importance, nicks to date have been less of a focus for study than DSBs. Previously available methods offered low resolution, preventing the identification of sequence biases and genomic features that affect nick accumulation and repair. Here we demonstrated the capability of NickSeq to measure the off-target activity of Cas9 nickase. NickSeq could also be beneficial in other contexts where nick detection is relevant such as analysis of damage caused by reactive oxygen species and other genotoxic agents or the relationship between nick accumulation and neurodegeneration in diseases characterized by deficient nick repair, telomere shortening, and cell death. Furthermore, as recently reported by Cao, et al.,^29^ nick detection methods like NickSeq could be applied to study other types of DNA damage as well by converting the damage into nicks; for example, base modifications could be studied through treatment with a glycosylase and AP endonuclease from the base excision repair pathway prior to readout by NickSeq.

## MATERIAL AND METHODS

### Preparation of DNA containing nicks

Biotinylated double-stranded oligonucleotides with one nick each were generated by annealing single-stranded oligonucleotides ordered from Integrated DNA Technologies. Nine oligos were ordered in total: a constant bottom strand (89 nucleotides long) used for each annealing reaction, four 5’ top strands of varying lengths (54, 55, 56, and 57 nucleotides) with 5’ biotin-TEG modifications, and four 3’ top strands of varying lengths (35, 34, 33, and 32 nucleotides) (Fig. 2a). Annealing was performed by creating an equimolar mixture of the bottom strand and a top strand pair, denaturing any base pairing by heating to 95 °C on a thermocycler, and reducing the temperature to 25 °C at a rate of 0.1 °C per second.

Nicking endonuclease Nb.BsmI (New England Biolabs) was used to incorporate nicks in one of the plasmids (extracted from bacteria with the Plasmid Plus Midi Kit, Qiagen) used. Two nicks, both on the reference strand, were expected based on sequence. A 10 μL reaction was prepared containing 1x NEBuffer 3.1 (New England Biolabs), 5 units of endonuclease, and 100 ng of plasmid and incubated at 65 °C for one hour. Enzyme inactivation then occurred by incubation at 80 °C for 20 minutes.

EnGen Spy Cas9 nickase (New England Biolabs), a Cas9 nuclease variant containing a D10A mutation in the RuvC nuclease domain, was used in conjunction with guide RNA (Synthego, synthetic cr:tracrRNA kit) to incorporate nicks in the other plasmid used as well as the *E. coli* genomic DNA (Thermo Scientific; genomic DNA purified from *E. coli* type B cells, ATCC 11303 strain) and human genomic DNA (extracted from cells with DNeasy Blood & Tissue Kit, Qiagen). One guide was used with the plasmid, expected to produce one nick on the non-reference strand. Two guides were used with the *E. coli* genome, expected to produce eight and one nicks respectively based on sequence. One guide, targeting the AAVS1 locus on chromosome 19, was used with the human genome. crRNA and tracrRNA were rehydrated and annealed according to the manufacturer’s instructions. For plasmid and *E. coli* genomes, 30 μL reactions were prepared containing 1x NEBuffer 3.1, 50 pmol of each guide RNA being used, 2 pmol Cas9 nickase, and 300 ng of DNA and incubated at 37 °C for an hour. Enzyme inactivation then occurred by incubation at 65 °C for 5 minutes. For reactions with human genomic DNA, a larger reaction volume and 10 μg of DNA were used.

### Incorporation of degenerate and biotinylated nucleotides

The degenerate nucleotides dPTP (TriLink Biotechnologies) and dKTP (Axxora) were incorporated into DNA at nicks via nick translation with *Taq* DNA polymerase (New England Biolabs). A 10 μL reaction was prepared containing 1x ThermoPol buffer (New England Biolabs), 0.25 units of polymerase, 20 pmol each of dPTP and dKTP, and 10 ng of DNA (or 10 μg of DNA in a larger reaction volume in the case of human genome) and incubated at 72 °C for 15 minutes. Polymerase and excess nucleotides were removed by 1:1 SPRI with Agencourt AMPure XP beads (Beckman Coulter) followed by elution in TE buffer.

Next, nick translation was performed with regular dNTPs (New England Biolabs) and biotin-11-dUTP (Thermo Scientific). Optimal results were obtained with biotinylated dUTP and dTTP in a 1:5 ratio. The purified DNA was incubated at 72°C for 30 minutes with 1x ThermoPol buffer, 0.25 units *Taq* polymerase, and 0.4 pmol of each dNTP (a 1 in 5,000 dilution compared to the manufacturer’s recommended PCR protocol) plus biotinylated dUTP. Polymerase and excess nucleotides were again removed by 1:1 SPRI and elution in TE buffer. During experiments with the oligonucleotides only (Fig. 2), biotinylated dUTP was not used during the second nick translation (since the oligonucleotides already contained a 5’ biotin-TEG modification). During experiments with human genomic DNA only, desthiobiotin-7-dATP (Jena Bioscience) was used in place of biotinylated dUTP for compatibility with the library preparation protocol.

### Targeted pulldown and library construction

Plasmid DNA and bacterial genome were tagmented at 55 °C for 10 minutes in a 20 μL reaction containing 5 mM magnesium chloride, 10 mM tris acetate, and 4 μL TDE1 (tagment DNA enzyme, Illumina). 0.1% SDS treatment then denatured the enzyme and 1:1 SPRI followed by elution in TE buffer was used to remove SDS. Human genomic DNA was fragmented by a dsDNA fragmentase (New England Biolabs) followed by end preparation and adapter ligation according to the NEBNext Ultra II DNA Library Prep Method for Illumina (New England Biolabs).

25 μg of Dynabeads MyOne Streptavidin C1 (Invitrogen) were washed in 100 μL of bind and wash buffer, 1 M sodium chloride and 0.1% tween-20 in TE buffer. DNA was diluted to 150 μL in bind and wash buffer and incubated on the beads for 30 minutes at room temperature with occasional gentle manual agitation. Following 3 washes in 180 μL bind and wash buffer, beads and bound biotinylated DNA were resuspended in 20 μL of TE buffer plus 0.01% tween-20. In experiments with the oligonucleotides only (Fig. 2), treatment with 100 mM sodium hydroxide for 5 minutes was used after the third wash followed by a final wash and resuspension to disrupt base pairing and allow for purification of the biotinylated strand (which initially contained the nick) only. In experiments with human genomic DNA only, the final wash was followed by a 20 minute incubation in 10 μL of 0.11 mg/mL biotin (Sigma-Aldrich) to elute desthiobiotinylated DNA from the beads.

1 μL of DNA (with beads unless elution was performed) was the substrate for a 10 μL qPCR reaction using either *Taq* DNA polymerase or KAPA HiFi (KAPA HiFi HotStart PCR kit, Kapa Biosystems) according to the manufacturer’s protocol with barcoded P5 and P7 sequencing primers at 0.5 mM and with 1x EvaGreen dye (Biotium) and 1x ROX reference dye (Invitrogen) used to monitor DNA amplification. Reagents and short primer-derived products were removed by 1:1 SPRI followed by elution in TE buffer.

### Sequencing

Enriched libraries were quantified using the Qubit dsDNA high sensitivity assay kit (Invitrogen) according to the manufacturer’s instructions. DNA was diluted to 1 nM and prepared for paired-end sequencing on a MiniSeq (Illumina) using a 75 or 150-cycle kit according to the manufacturer’s instructions with a final loading concentration of 1.1 pM.

Following sequencing and barcode-based demultiplexing, Cutadapt^50^ removed primer sequences present from both the 5’ and 3’ ends of the reads. Bowtie2^51^ aligned reads to the appropriate reference genome, discarding read pairs that mapped discordantly. Experiments on human genomic DNA were mapped to GRCh37 (hg19) in order to exclude reads mapping to ENCODE blacklist regions from analysis and compare results with literature that also mapped to this reference. SAMtools^52^ created two sorted and indexed *.bam files: one including all reads and one with duplicate reads removed. Peaks in sequencing coverage were identified by MACS2^33^, a peak caller originally created for use with ChIP-seq data.

### Computational analysis

Further analysis of sequencing data was performed with custom scripts written in Python 3.6.8. Identification of nicks genome-wide began by assigning a q-value to all nucleotides within MACS2 peaks with q-scores greater than 0.4 (this inclusion of low-quality peaks means many false-positive peaks occur, but use of the mutational signal prevents false-positive nick calls in these peaks). Q-values were based on binomial tests determining the probability of receiving the observed transition mutation rate (C↔T and A↔G, the only mutations caused by dPTP and dKTP incorporation) at each location with the null expectation being that the observed rate is equal to the maximum of either the observed transversion (non-transition) mutation at that location or 0.03. Potential nicks were determined as locations with sufficiently low q-values, as this would suggest the observed transition mutation rate at that location is due to dPTP or dKTP incorporation and not to other factors such as mis-incorporation during PCR, sequencing errors, or mapping errors. These sites were then filtered based on proximity to one another; that is if a site did not have another site within close proximity (less than five nucleotides) it was removed as dPTP and dKTP incorporation is expected to produce of string of sequential mutations under the conditions of our protocol. Final filtering to remove locations with very low read count yields the final list of predicted nick loci.

At this point, each locus with a predicted nick consists of the range of locations over which there was mutational signal with the nick occurring on one end of the range. Comparing the nick location to the location of maximum coverage in the MACS2 peak reveals which end of the range as well as which strand of DNA the nick likely occurred on. Since DNA is always synthesized 5’→3’, the location of maximum coverage is expected 3’ relative to the nick. Therefore, if the nick occurs 5’ to the peak’s maximum on the reference strand it is a reference strand nick at the 5’ end of the predicted range. If the nick occurs 3’ to the peak’s maximum on the reference strand (which is 5’ on the non-reference strand) it is a non-reference strand nick at the reference strand’s 3’ end (non-reference strand’s 5’ end) of the predicted range. During this analysis and in all figures, mutation rates are determined from the *.bam file with all sequencing reads and sequencing coverage is determined from the *.bam with duplicate reads removed (using the *.bam with all sequencing reads slightly alters figure appearance but does not alter nick prediction).

## ACKNOWLEDGEMENT

The authors thank Arnaud Gutierrez for discussions in designing the NickSeq method as well as Dr. David Liu and members of his laboratory for discussions about the application of NickSeq to genome editing. We also thank all members of the Blainey laboratory for discussions and providing materials used in this work.

---

**Figure S1:**
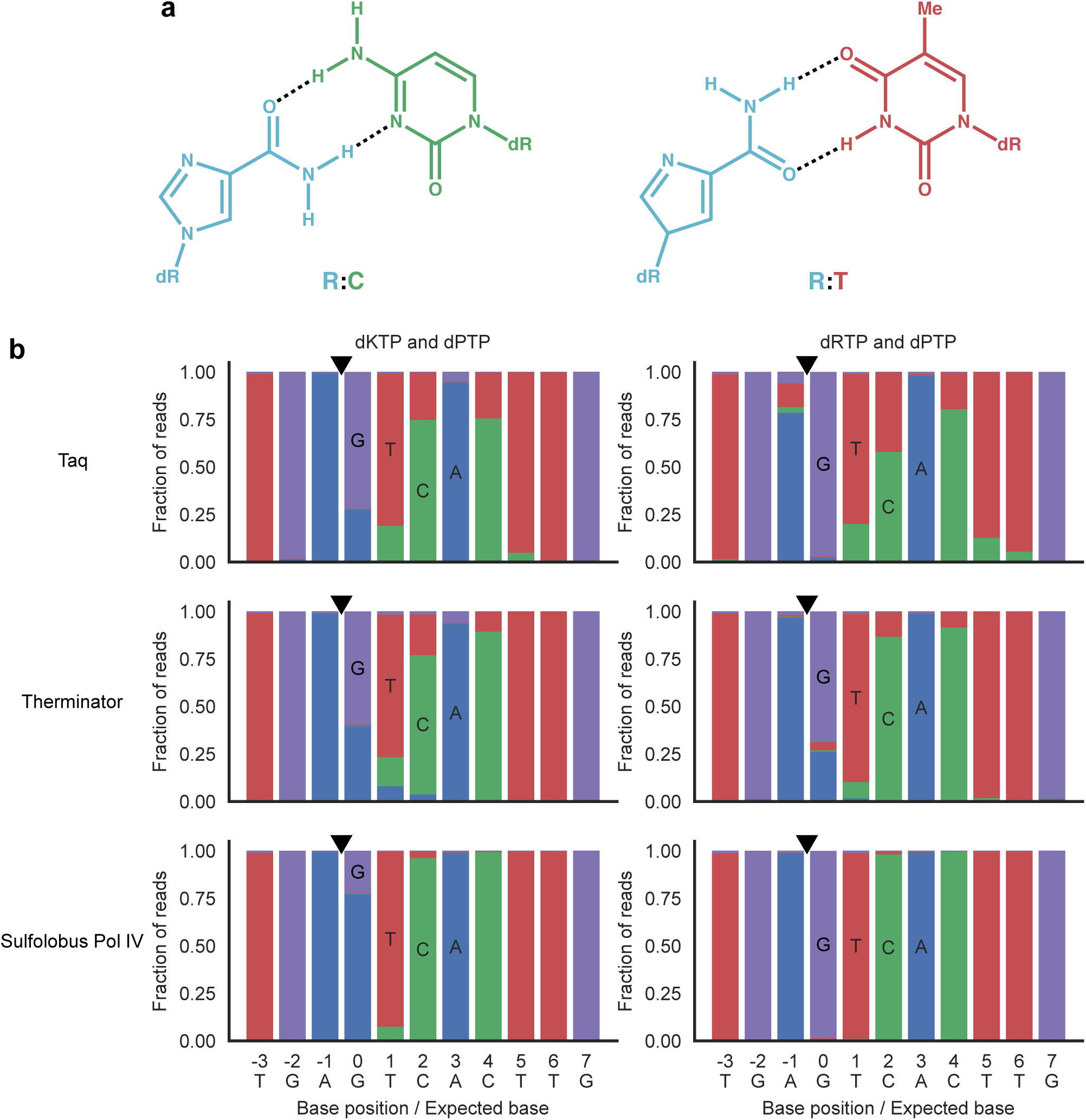
Effect of dRTP during PCR. **a)** Ribavirin (RTP) is a nucleotide that displays antiviral activity. Due to its structure, we considered deoxyribavirin (dRTP) as an alternative to dKTP when performing NickSeq for the ability to base pair with both C and T residues. Dashed black lines represent hydrogen bonds. **b)** dKTP and dRTP (Biolog Life Science Institute, special order) were compared for their mutagenic abilities by incorporation into the nick-containing biotinylated oligonucleotide (Fig. 2a). Initial nick translations were performed with either *Taq* DNA polymerase, Therminator DNA polymerase (New England Biolabs), or *Sulfolobus* DNA polymerase IV (New England Biolabs). *Taq* and dRTP lead to an unexpected result: a mutational signal 5’ of the SSB that would interfere with precise resolution of SSB location. Pol IV led to a stronger signal one nucleotide 3’ of the SSB with dKTP but much weaker signal at consecutive nucleotides and was unable to incorporate dRTP. While Therminator would incorporate dRTP, it does not appear to provide any significant benefit over using *Taq* with dKTP.

**Figure S2:**
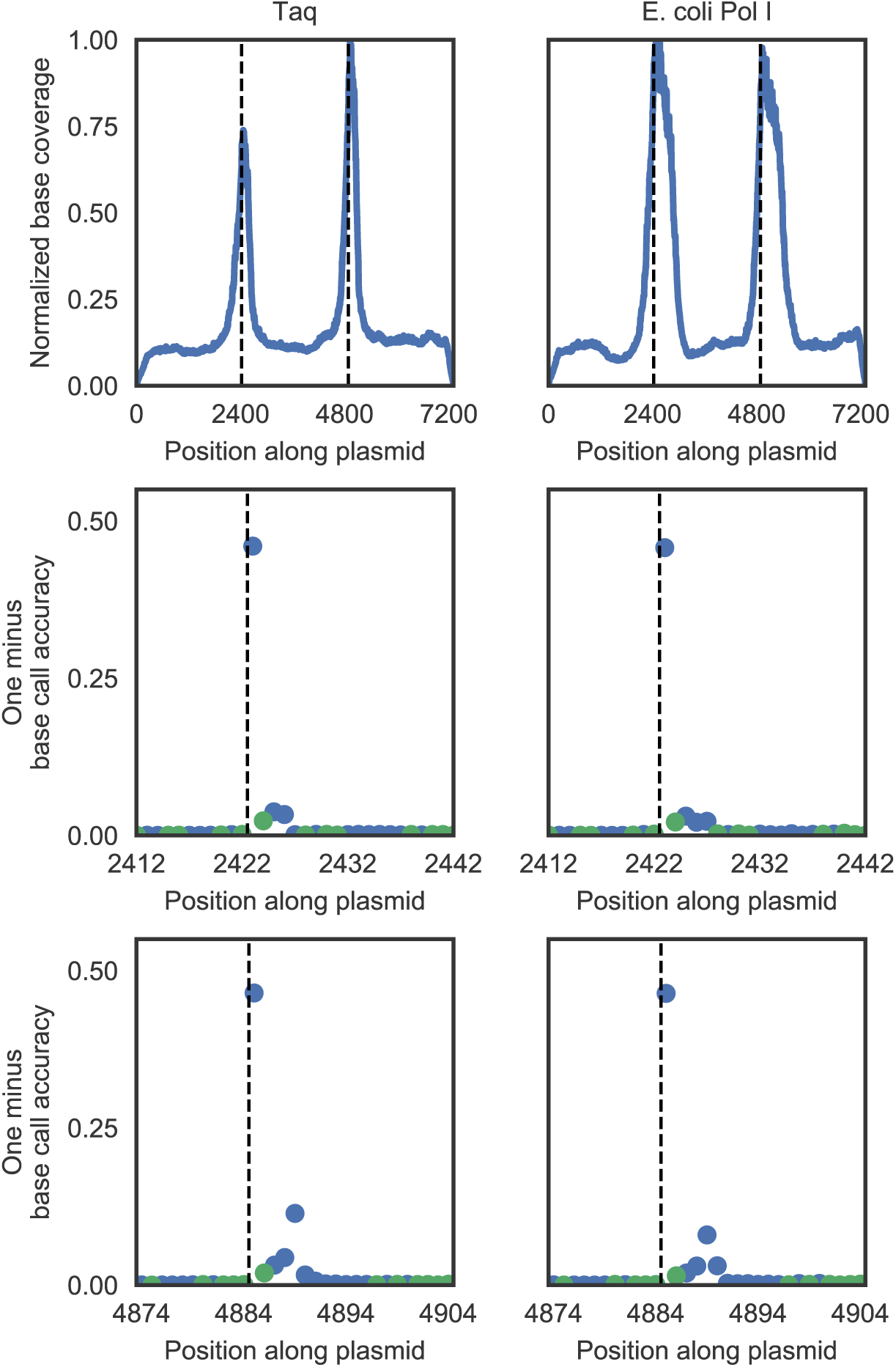
Different DNA polymerases can be used during the second nick translation. *E. coli* DNA polymerase I (New England Biolabs) can be used in place of *Taq* DNA polymerase during nick translation with dNTPs and biotin-dUTP. Sequencing peaks are wider with Pol I when nearly identical protocols are used (the only difference being Pol I incubates at 37 °C), however if this is undesirable a shorter incubation time will result in more narrow peaks. Use of Pol I could potentially lead to lower background noise since, unlike *Taq*, it has 3’→5’ exonuclease ‘proofreading’ activity that results in a lower observed base mis-incorporation rate.

**Figure S3:**
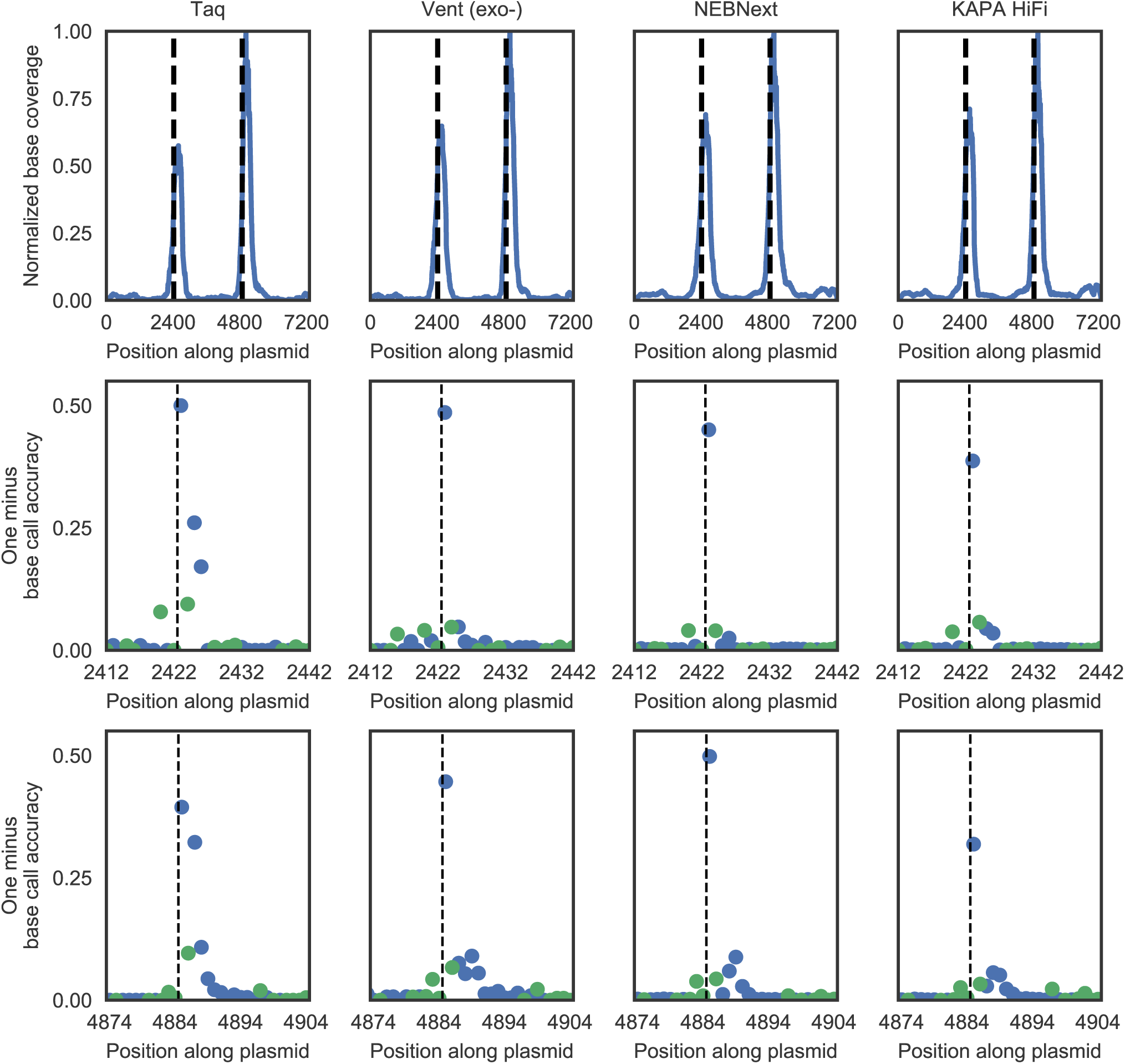
Different DNA polymerases can be used during PCR. In addition to *Taq* DNA polymerase and KAPA HiFi, Vent (exo-) DNA polymerase (New England Biolabs) and NEBNext (New England Biolabs) can be used during PCR. Each enzyme results in slightly different signal intensities from the incorporated degenerate bases. However, while *Taq* has higher signal than KAPA HiFi, for example, it also has a higher level of background noise due to its higher base mis-incorporation rate.

**Figure S4:**
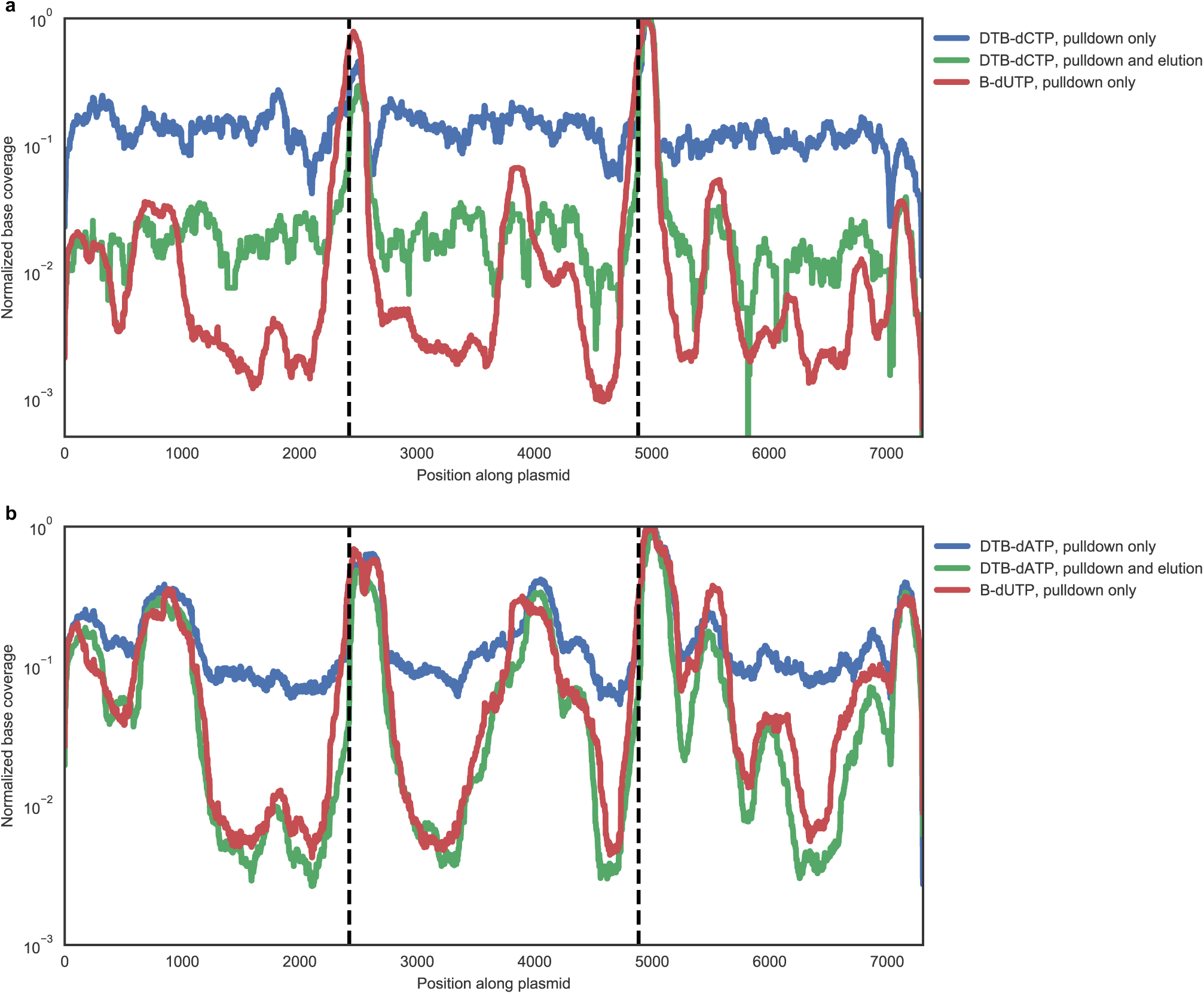
Desthiobiotinylated nucleotides provide an alternative purification method involving elution from the streptavidin beads. At high concentrations, the magnetic streptavidin beads used for library enrichment can inhibit PCR. To overcome this, desthiobiotinylated nucleotides that could be eluted from the beads were tested as an alternative to biotinylated nucleotides. Desthiobiotin is an elutable biotin derivative with reduced affinity for streptavidin. Desthiobiotinylated DNA was eluted by resuspending beads in 10 μL of 0.11 mg/mL biotin in water. After 20 minutes, supernatant with desthiobiotinylated DNA was collected from the beads. **a)** Desthiobiotin-aminoallyl-dCTP (TriLink Biotechnologies) did not provide as good library enrichment as biotin-dUTP. **b)** Desthiobiotin-aminohexyl-dATP (Jena Bioscience) provided slightly better library enrichment than biotin-dUTP.

**Figure S5:**
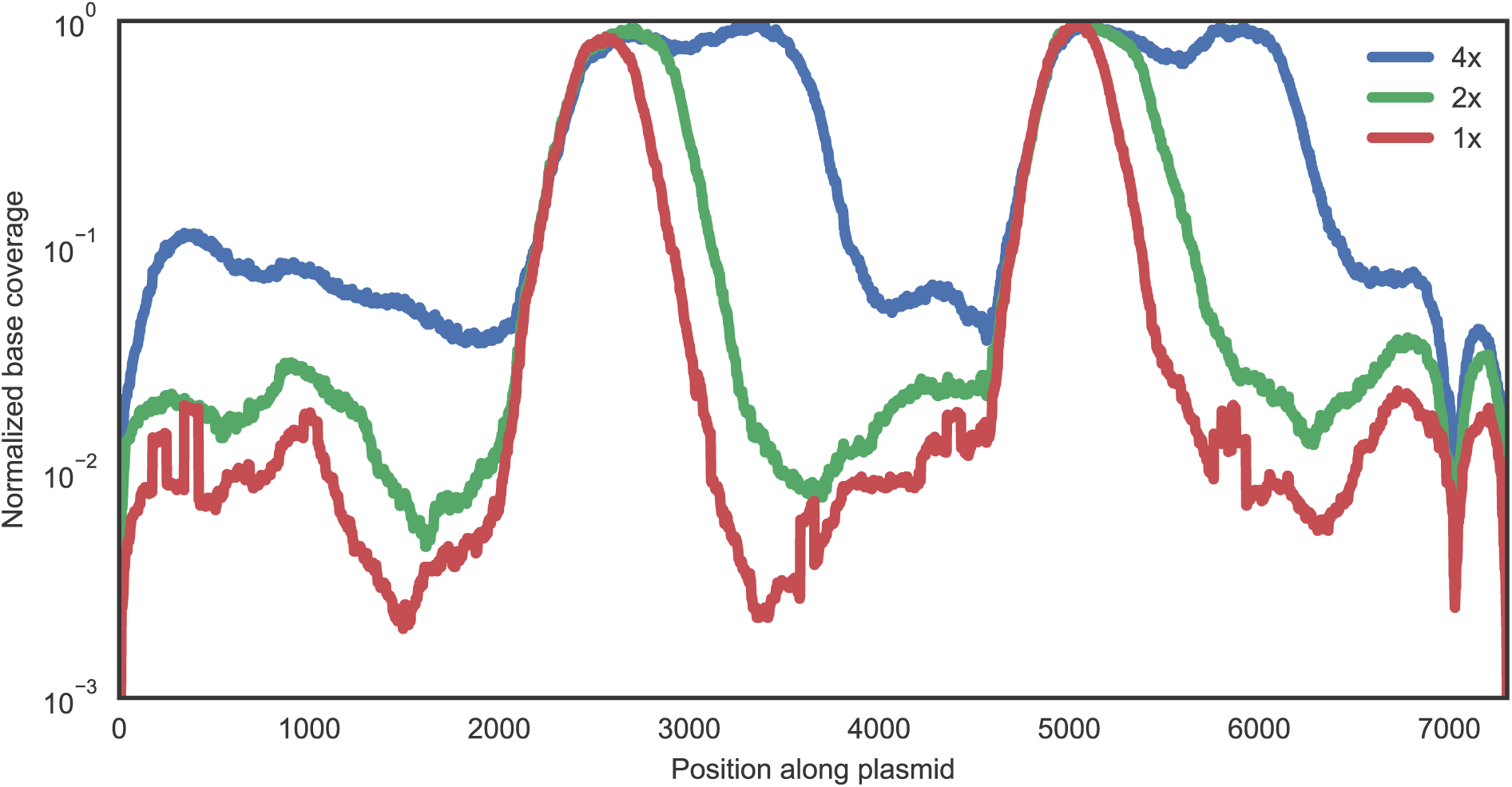
Sequencing coverage peaks widen with increasing dNTP and biotinylated dUTP concentrations used during nick translation. The optimal concentration of dNTPs during the second nick translation was determined to be 40 nM with a 1:5 ratio of biotinylated dUTP to dTTP (defined as 1x). At higher concentrations, the peaks in sequencing coverage around the nicks become wider and there is additional background at locations far from nicks.

**Figure S6:**
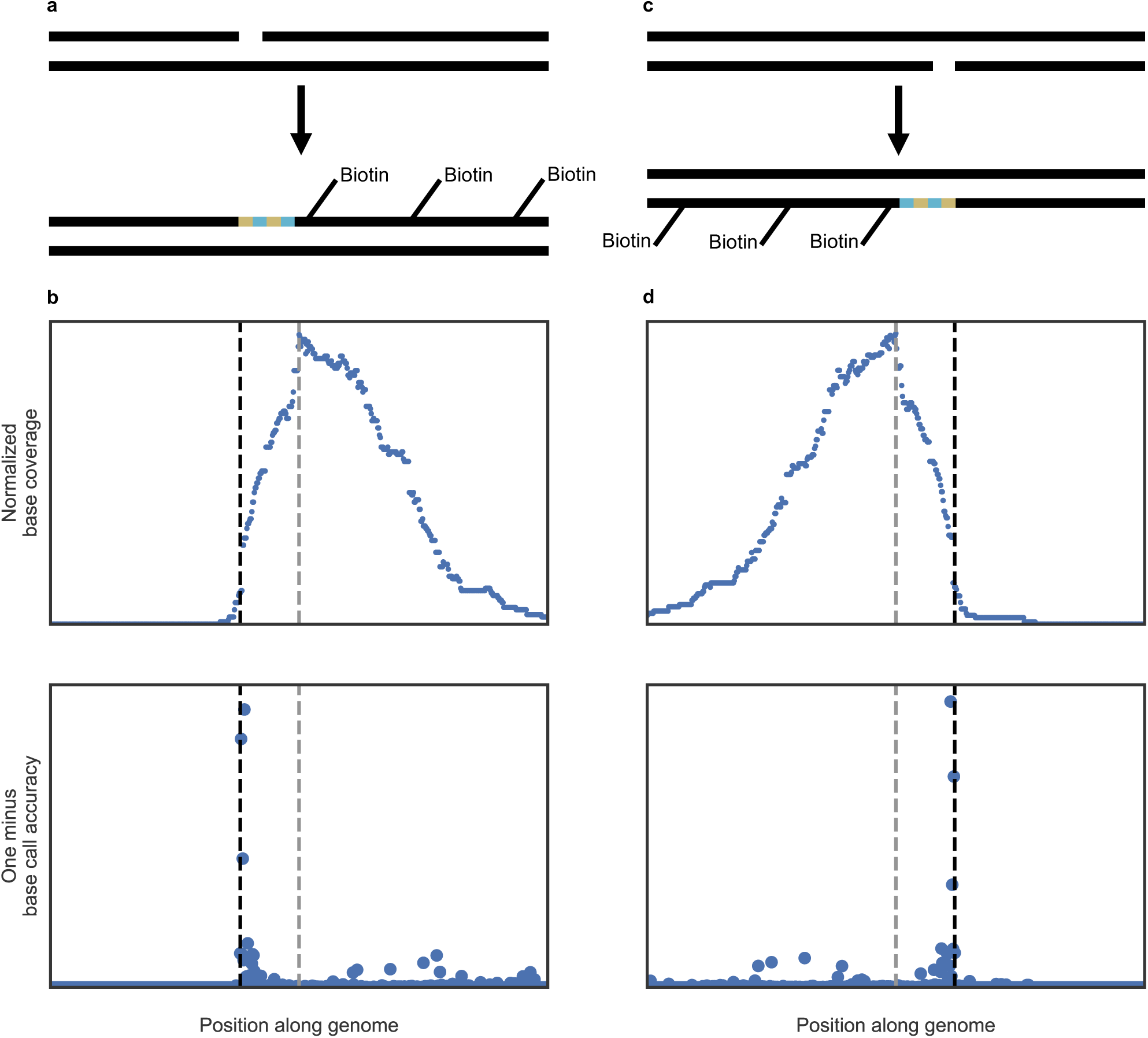
Relative locations of the mutational signal and peak in sequencing coverage reveal the nicked strand. DNA is always synthesized in the 5’→3’ direction. Therefore, the biotinylation appears either 3’ (**a**) or 5’ (**c**) of the incorporated P and K residues with respect to the reference strand depending on the strand the nick was present in. Upon sequencing, the peak in coverage around the nick (gray dashed line) will appear downstream (**b**) or upstream (**d**) of the mutational signal (black dashed line) if the nick was on the reference or non-reference strand, respectively.

**Figure S7:**
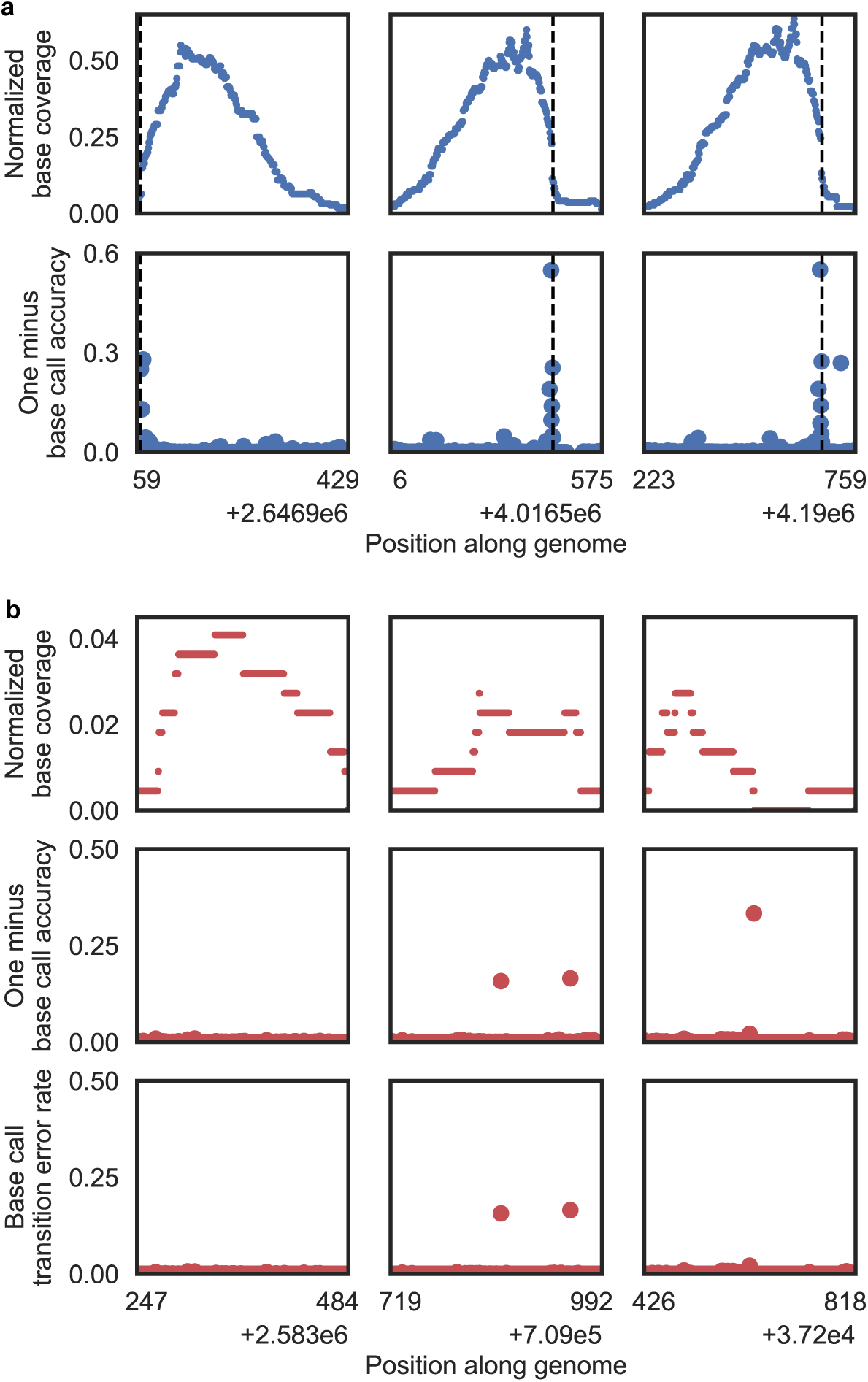
Representative sequencing peaks in the *E. coli* genome called by MACS2 that did and did not contain nicks. **a)** In total, nine nicks were expected to be incorporated into the *E. coli* genome as a result of D10A spCas9 nickase and the two guide RNAs used. One guide RNA had a single target site (Fig. 5d), while the other had eight, five of which are shown in Fig. 5c and the rest here. **b)** Nicks with low penetrance in a sample cannot be identified from sequencing coverage and peak calling alone as background peaks may result from non-specific binding of DNA to the streptavidin beads used for purification. The mutational signal that is unique to NickSeq can be used to filter out peaks that do not contain nicks, allowing for detection of such low penetrance nicks. Other metrics such as peak height or quality and p-values cannot recapitulate this level of performance (Fig. 5b). Sequencing peaks called by MACS2 may have very little mutational signal (left). Alternatively, signal may be present but not occur at consecutive loci as would be expected from incorporation of consecutive P and K residues (middle). Finally, mutational signal may be detected that does not represent a transition mutation (C↔T or A↔G) and therefore could not have resulted from P and K residues being incorporated (right). The latter two scenarios can occur from real variants in the sample DNA relative to the reference, misincorporations during PCR, or sequencing errors.

**Table S1:**
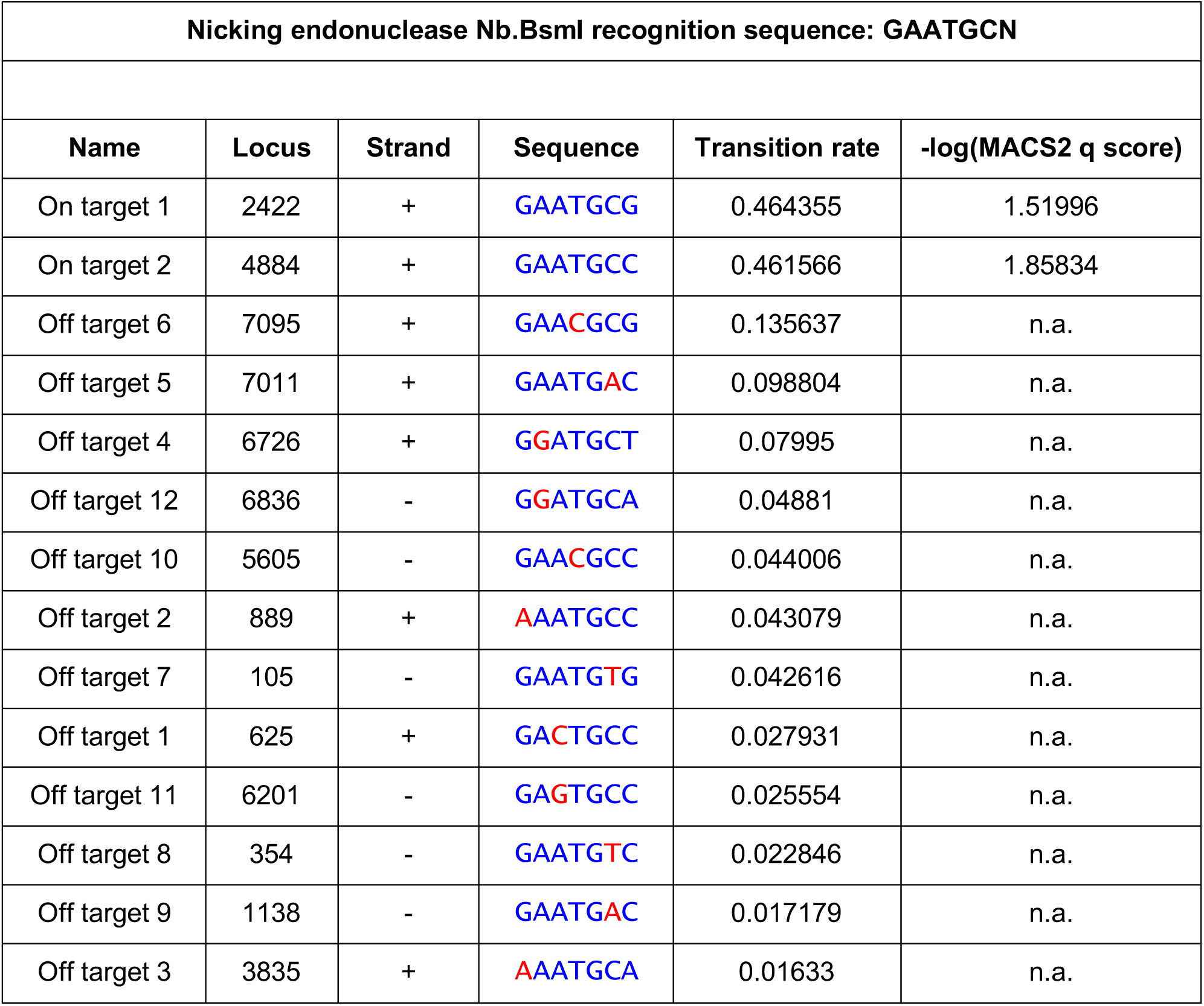
Off-target nicks detected after treatment of a plasmid with nicking endonuclease Nb.BsmI.

**Table S2:**
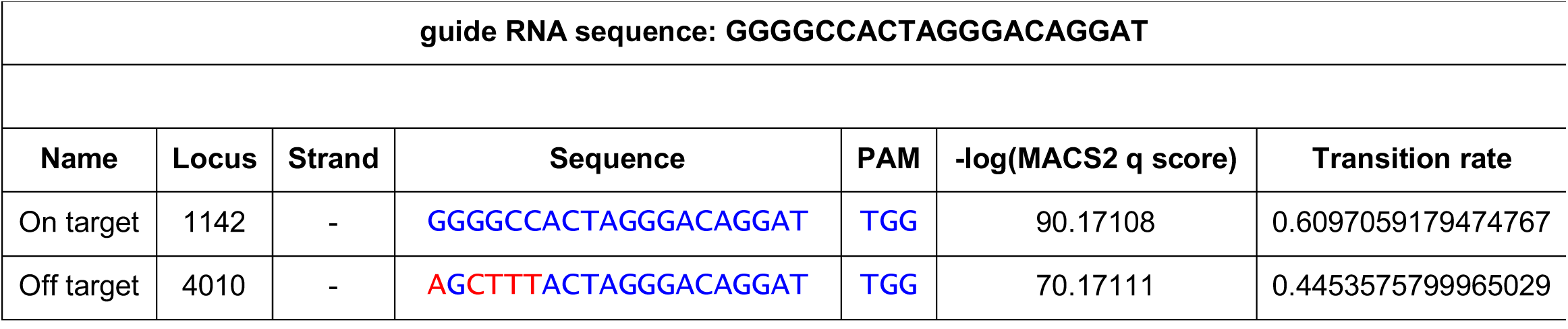
Nicks detected after treatment of a plasmid with D10A spCas9 nickase.

**Table S3:**
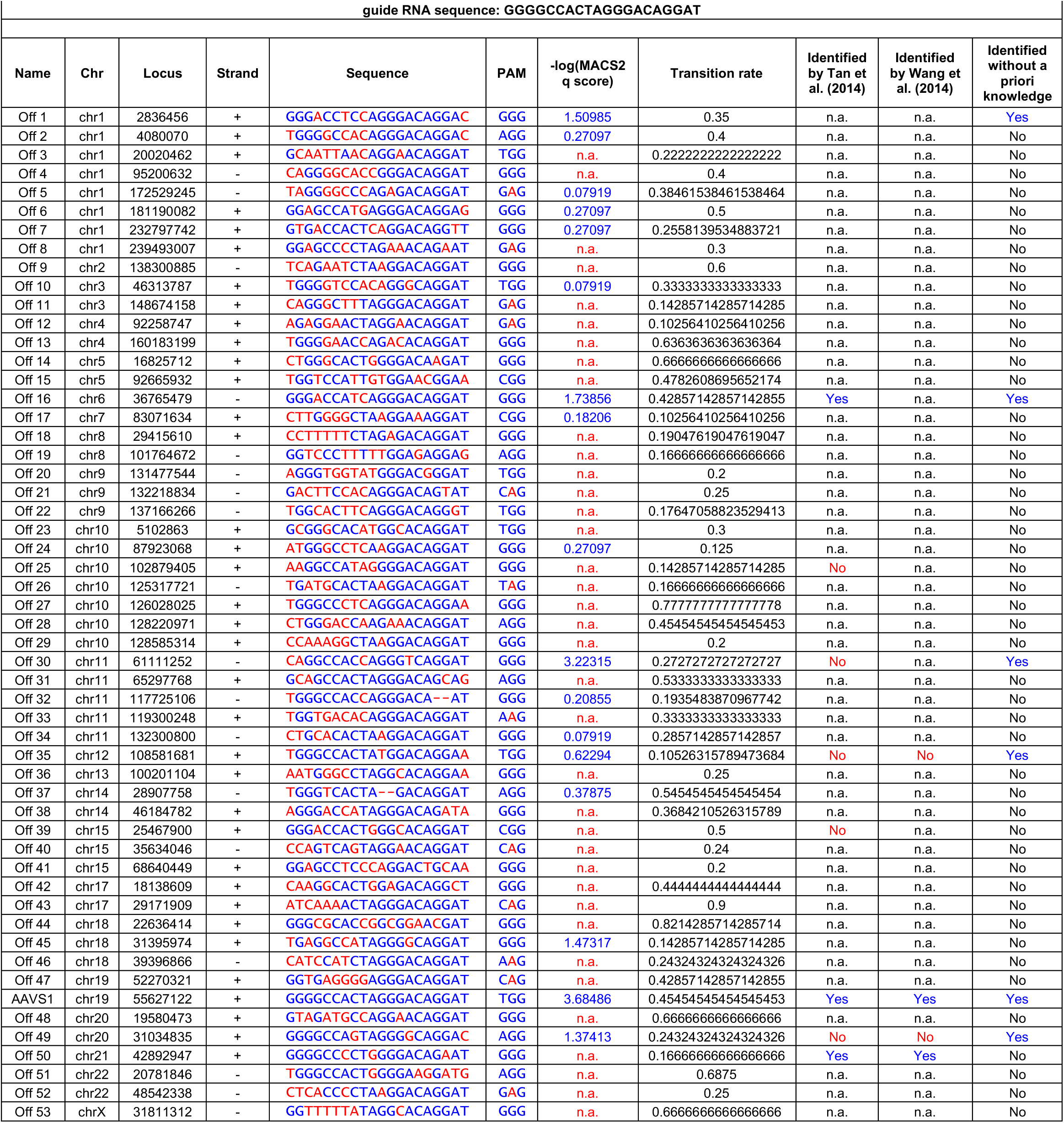
Detected off-target activity of D10A spCas9 nickase on human genomic DNA.

